# VSGs expressed during natural *T. b. gambiense* infection exhibit extensive sequence divergence and a subspecies-specific expression bias

**DOI:** 10.1101/2021.09.09.459620

**Authors:** Jaime So, Sarah Sudlow, Abeer Sayeed, Tanner Grudda, Stijn Deborggraeve, Dieudonné Mumba Ngoyi, Didier Kashiama Desamber, Bill Wickstead, Veerle Lejon, Monica R. Mugnier

## Abstract

*Trypanosoma brucei gambiense* is the primary causative agent of human African trypanosomiasis (HAT), a vector-borne disease endemic to West and Central Africa. The extracellular parasite evades antibody recognition within the host bloodstream by altering its Variant Surface Glycoprotein (VSG) coat through a process of antigenic variation. The serological tests which are widely used to screen for HAT use VSG as one of the target antigens. However, the *VSGs* expressed during human infection have not been characterized. Here we use VSG-seq to analyze the *VSGs* expressed in the blood of patients infected with *T. b. gambiense* and compared them to VSG expression in *T. b. rhodesiense* infections in humans as well as *T. b. brucei* infections in mice. The 44 *VSGs* expressed during *T. b. gambiense* infection revealed a striking bias towards expression of type B N-termini (82% of detected VSGs). This bias is specific to *T. b. gambiense*, which is unique among *T. brucei* subspecies in its chronic clinical presentation and anthroponotic nature, pointing towards a potential link between *VSG* expression and pathogenesis. The expressed *T. b. gambiense VSGs* also share very little similarity to sequences from 36 *T. b. gambiense* whole genome sequencing datasets, particularly in areas of the VSG protein exposed to host antibodies, suggesting that wild *T. brucei VSG* repertoires vary more than previously expected. Overall, this work demonstrates new features of antigenic variation in *T. brucei gambiense* and highlights the importance of understanding *VSG* repertoires in nature.

**Significance Statement:** Human African Trypanosomiasis is a neglected tropical disease primarily caused by the extracellular parasite *Trypanosoma brucei gambiense*. To avoid elimination by the host, these parasites repeatedly replace their Variant Surface Glycoprotein (VSG) coat. Despite the important role of VSGs in prolonging infection, *VSG* expression during human infections is poorly understood. A better understanding of natural *VSG* gene expression dynamics can clarify the mechanisms that *T. brucei* uses to alter its VSG coat and improve trypanosomiasis diagnosis in humans. We analyzed the expressed *VSGs* detected in the blood of patients with trypanosomiasis. Our findings indicate that there are features of antigenic variation unique to human-infective *T. brucei* subspecies and *VSGs* expressed in natural infection may vary more than previously expected.

## Introduction

Human African Trypanosomiasis (HAT) is caused by the protozoan parasite *Trypanosoma brucei. T. brucei* and its vector, the tsetse fly, are endemic to sub-Saharan Africa (1). There are two human-infective *T. brucei* subspecies: *T. b. gambiense*, which causes chronic infection in West and Central Africa (~98% of cases), and *T. b. rhodesiense*, which causes acute infection in East and Southern Africa (~2% of cases) (2, 3). In humans, infections progress from an early stage, usually marked by a fever and body aches, to a late stage associated with severe neurological symptoms that begins when the parasite crosses the blood-brain barrier (4). HAT is considered fatal without timely diagnosis and treatment. While around 50 million people are at risk of infection (5), the number of annual human infections has declined significantly in recent years, with only 864 cases reported in 2019 (6). The World Health Organization is working towards zero human transmissions of HAT caused by *T. b. gambiense* (gHAT) by 2030 (7). Case detection and treatment is an important component of current public health initiatives to control the disease.

Prospects for developing a vaccine are severely confounded by the ability of African trypanosomes to alter their surface antigens (8). As *T. brucei* persists extracellularly in blood, lymph, and tissue fluids, it is constantly exposed to host antibodies (9–12). The parasite periodically changes its dense Variant Surface Glycoprotein (VSG) coat to evade immune recognition. This process, called antigenic variation, relies on a vast collection of thousands of VSG-encoding genes (13–16). *T. brucei* also continually expands the number of usable antigens by constructing mosaic *VSGs* through one or more recombination events between individual *VSG* genes (17, 18).

Although the *VSG* repertoire is enormous and potentially expanding, these variable proteins are the primary antigens used for serological screening for gHAT (there is currently no serological test for diagnosis of infection with *T. b. rhodesiense*). One VSG in particular, LiTat 1.3, has been identified as an antigen against which many gHAT patients have antibodies (19) and thus serves as the main target antigen in the primary serological screening tool for gHAT, the card agglutination test for trypanosomiasis (CATT/*T. b. gambiense*) (20). More recently developed rapid diagnostic tests use a combination of native LiTat1.3 and another VSG, LiTat1.5 (21, 22), or the combination of a VSG with the invariant surface glycoprotein ISG 65 (23).

Despite the widespread use of VSGs as antigens to screen for gHAT, little is known about how the large genomic repertoire of *VSGs* is used in natural infections; the number and diversity of *VSGs* expressed by wild parasite populations remain unknown. It is unclear whether VSG repertoires are evolving in the field, potentially affecting the sensitivity of serological tests that use VSG as an antigen. Notably, some *T. b. gambiense* strains lack the LiTat 1.3 gene entirely (24, 25). A study from our lab that evaluated VSG expression during experimental mouse infections by VSG-seq, a targeted RNA-sequencing method that identifies the *VSGs* expressed in a given population of *T. brucei*, revealed significant *VSG* diversity within parasite populations in each animal (26). This diversity suggested that the parasite’s genomic VSG repertoire might be insufficient to sustain a chronic infection, highlighting the potential importance of the recombination mechanisms that form new VSGs (13, 17).

Given the role of VSGs during infection and their importance in gHAT screening tests, a better understanding of *VSG* expression in nature could inform the development of improved screening tests while providing insight into the molecular mechanisms of antigenic variation. To our knowledge, only one study has investigated *VSG* expression in wild *T. brucei* isolates (27). For technical reasons, this study relied on RNA isolated from parasites passaged through small animals after collection from the natural host. As *VSG* expression may change during passage, the data obtained from these samples are somewhat difficult to interpret. To better understand the characteristics of antigenic variation in natural *T. brucei* infections, we sought to analyze *VSG* expression in *T. brucei* field isolates from which RNA was directly extracted.

In the present study, we used VSG-seq to analyze the *VSGs* expressed by *T. b. gambiense* in the blood of 12 patients with a confirmed infection. To complement these data, we also used our pipeline to analyze published RNA-seq datasets from both experimental mouse infections and *T. b. rhodesiense* patients. In addition to VSG-seq, we searched for evidence of sequence homology in a large set of whole genome sequences for a variety of *T. b. gambiense* isolates. Our analysis revealed distinct biases in *VSG* expression that appear to be unique to the *T. b. gambiense* subspecies and a divergence between expressed patient *VSG* and previously characterized *T. b. gambiense* strains that suggests patient VSG repertoires are more diverse than previously expected.

## Results

### Parasites in gHAT patients express diverse sets of VSGs

To investigate *VSG* expression in natural human infections, we performed VSG-seq on RNA extracted from whole blood collected from 12 human African trypanosomiasis patients from five locations in the Kwilu province of the Democratic Republic of the Congo (DRC) (Figure 1A). We estimated the relative parasitemia of each patient by SL-QPCR (28), and we estimated the number of parasites after mAECT on buffy coat for all patients except patient 29 (Table 1). Using RNA extracted from 2.5 mL of whole blood from each patient, we amplified *T. brucei* RNA from host/parasite total RNA using a primer against the *T. brucei* spliced leader sequence and an anchored oligo-dT primer. The resulting trypanosome-enriched cDNA was used as a template to amplify *VSG* cDNA in three replicate reactions, and *VSG* amplicons were then submitted to VSG-seq sequencing and analysis. To determine whether a *VSG* was expressed within a patient, we applied the following stringent cutoffs:

1. We conservatively estimate that each 2.5 mL patient blood sample contained a minimum of 100 parasites. At this minimum parasitemia, a single parasite would represent 1% of the population (and consequently ~1% of the parasite RNA in a sample). As a result, we excluded all *VSGs* comprising <1% of the total VSG-seq pool in each patient as unlikely to represent the major expressed VSG in at least one cell from the population.
2. We classified *VSGs* as expressed if they met the expression cutoff in at least two of three technical library replicates.

**Figure 1.**
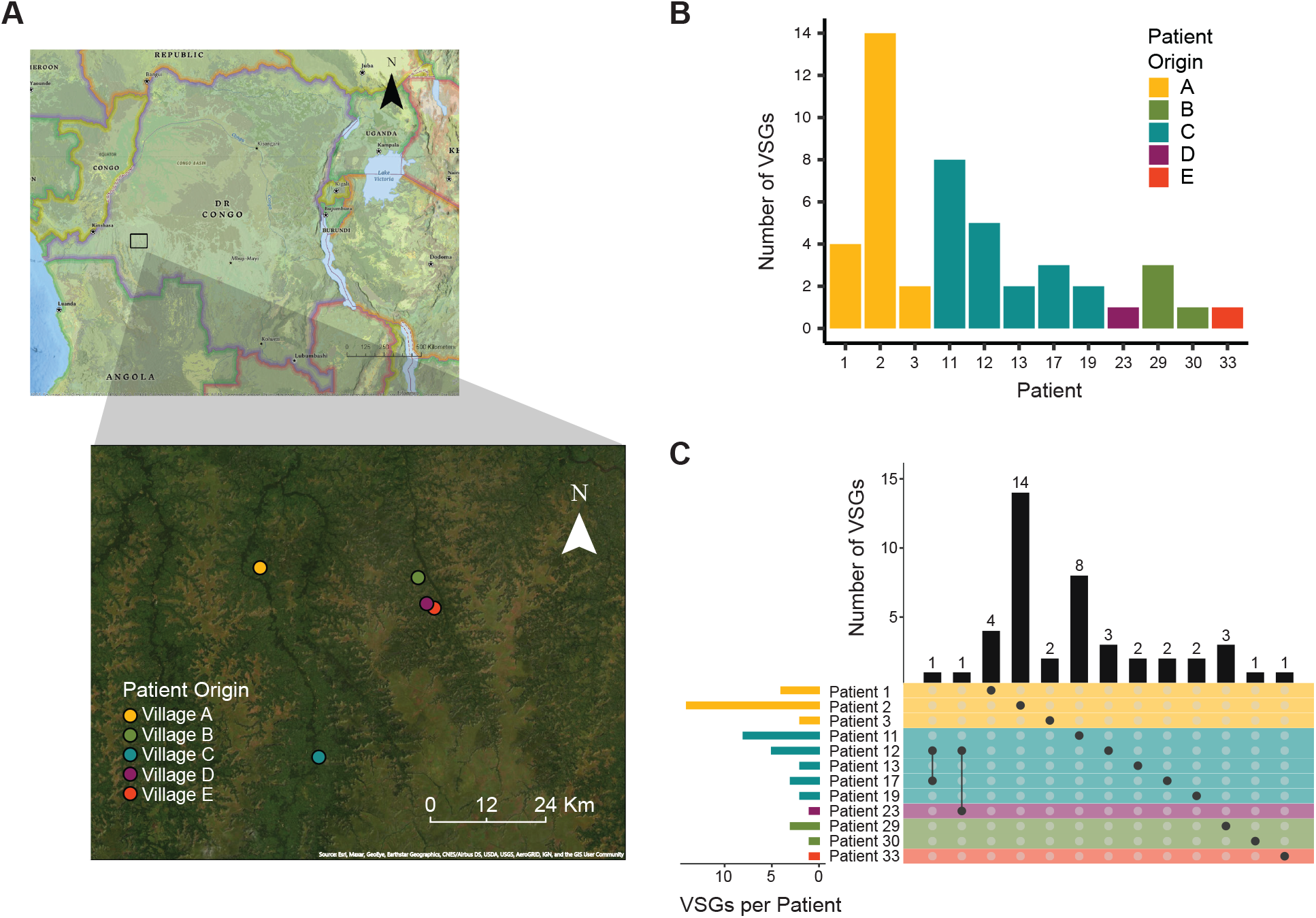
Parasites isolated from gHAT patients express multiple *VSGs*. (A) Map showing the location of each patient’s home village. Maps were generated with ArcGIS^®^ software by Esri, using world imagery and National Geographic style basemaps. (B) Graph depicting the total number of *VSGs* expressed in each patient. (C) The intersection of expressed *VSG* sets in each patient. Bars on the left represent the size of the total set of VSGs expressed in each patient. Dots represent an intersection of sets with bars above the dots representing the size of the intersection. Color indicates patient origin.

**Table 1.**
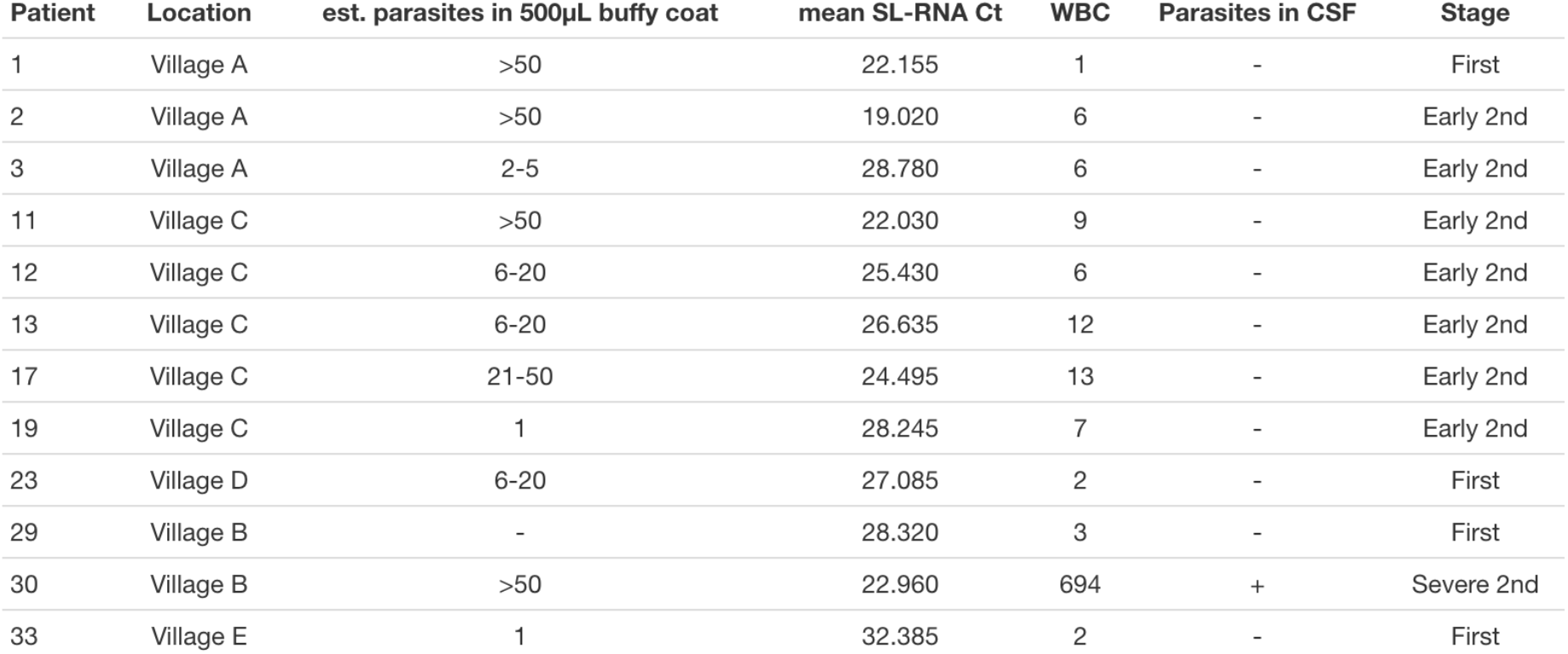
Patient stage and parasitemia data. We used the following staging definitions: First: 0-5 WBC/μl, no trypanosomes in cerebrospinal fluid (CSF). Second: >5 WBC/μl or trypanosomes in CSF (with early 2^nd^: 6-20 WBC/μl and no trypanosomes in CSF; severe 2^nd^: >100 WBC/μl). WBC: white blood cells.

| Patient | Location | est. parasites in 500µL buffy coat | mean SL-RNA Ct | WBC | Parasites in CSF | Stage |
| --- | --- | --- | --- | --- | --- | --- |
| 1 | Village A | >50 | 22.155 | 1 | - | First |
| 2 | Village A | >50 | 19.020 | 6 | - | Early 2nd |
| 3 | Village A | 2-5 | 28.780 | 6 | - | Early 2nd |
| 11 | Village C | >50 | 22.030 | 9 | - | Early 2nd |
| 12 | Village C | 6-20 | 25.430 | 6 | - | Early 2nd |
| 13 | Village C | 6-20 | 26.635 | 12 | - | Early 2nd |
| 17 | Village C | 21-50 | 24.495 | 13 | - | Early 2nd |
| 19 | Village C | 1 | 28.245 | 7 | - | Early 2nd |
| 23 | Village D | 6-20 | 27.085 | 2 | - | First |
| 29 | Village B | - | 28.320 | 3 | - | First |
| 30 | Village B | >50 | 22.960 | 694 | + | Severe 2nd |
| 33 | Village E | 1 | 32.385 | 2 | - | First |

1112 unique *VSG* open reading frames were assembled *de novo* from the patient reads and 44 met our expression criteria. Only these 44 *VSGs*, which we will refer to as “expressed *VSGs*,” were considered in downstream analysis, except when otherwise noted. TgsGP, the VSG-like protein which partially enables resistance to human serum in *T. b. gambiense* (29), assembled in samples from patients 2, 11, 13, and 17, and met the expression threshold in patients 2, 11, and 17. The absence of this transcript in most samples is likely due to the low amount of input material used to prepare samples.

At least one *VSG* met our expression criteria in each patient, and in most cases, multiple *VSGs* were detected. Patient 2 showed the highest diversity, with 14 *VSGs* expressed (Figure 1B, Supplemental Figure 1). There is a positive correlation between parasitemia, as estimated by qPCR, and the number of detected *VSGs* (Supplemental Figure 2), suggesting that Our blood volumes may not be sampling the full diversity of circulating expressed VSG at low parasitemia. Nevertheless, two *VSGs* were shared between patients: *VSG* ‘Gambiense 195’ was expressed in both patient 12 and patient 17 from Village C; *VSG*‘Gambiense 38’ was expressed in patient 12 from Village C and patient 23 from Village D (Figure 1C). Because our sampling did not reach saturation, resulting in some variability between technical replicates, we focused only on the presence/absence of individual VSGs for further analysis, rather than relative expression levels within each population.

### Natural *T. b. gambiense* infections show a strong bias towards the expression of type B VSG

To further characterize the set of expressed *VSGs* in these samples, we sought to define the VSG domain types encoded by each *VSG. T. brucei* VSG contains two domains: a variable N-terminal domain of ~350-400 amino acids, and a less variable C-terminal domain of ~40-80 amino acids, characterized by one or two conserved groups of four disulfide-bonded cysteines (13, 30). On the surface of trypanosomes, the VSG N-terminal domain is readily exposed to the host. In contrast, the C-terminal domain is proximal to the plasma membrane and largely hidden from host antibodies (31–33). The N-terminal domain is classified into two types, A and B, each further distinguished into subtypes (A1-3 and B1-2), while the C-terminal domain has been classified into six types (1–6) (13, 30). These classifications are based on protein sequence patterns anchored by the conservation of cysteine residues, but the biological implications of VSG domain types have not been investigated.

We evaluated two automated approached for determining the type and subtype of each VSG’s N-terminal domain. The first approach was to create a bioinformatic pipeline to determine each N-terminal domain subtype, using HMM profiles we created for each subtype from sets of N-terminal domains previously typed by Cross et al. (15). The second approach was to create a BLASTp network graph based on a published method (34) where the N-terminal subtype of a VSG is determined by the set of VSGs it clusters with, and clusters are identified using the leading eigenvector method (35). We used each approach to determine the N-terminal subtype of each expressed *VSG* in our patient sample dataset, along with 863 VSG N-termini from the Lister 427 genome. We compared these results to either existing N-terminal classification (for Lister 427 VSGs) or classification based on position in a newly-generated BLASTp-tree (15) (for *T. b. gambiense* VSGs; Figure 2A).

**Figure 2.**
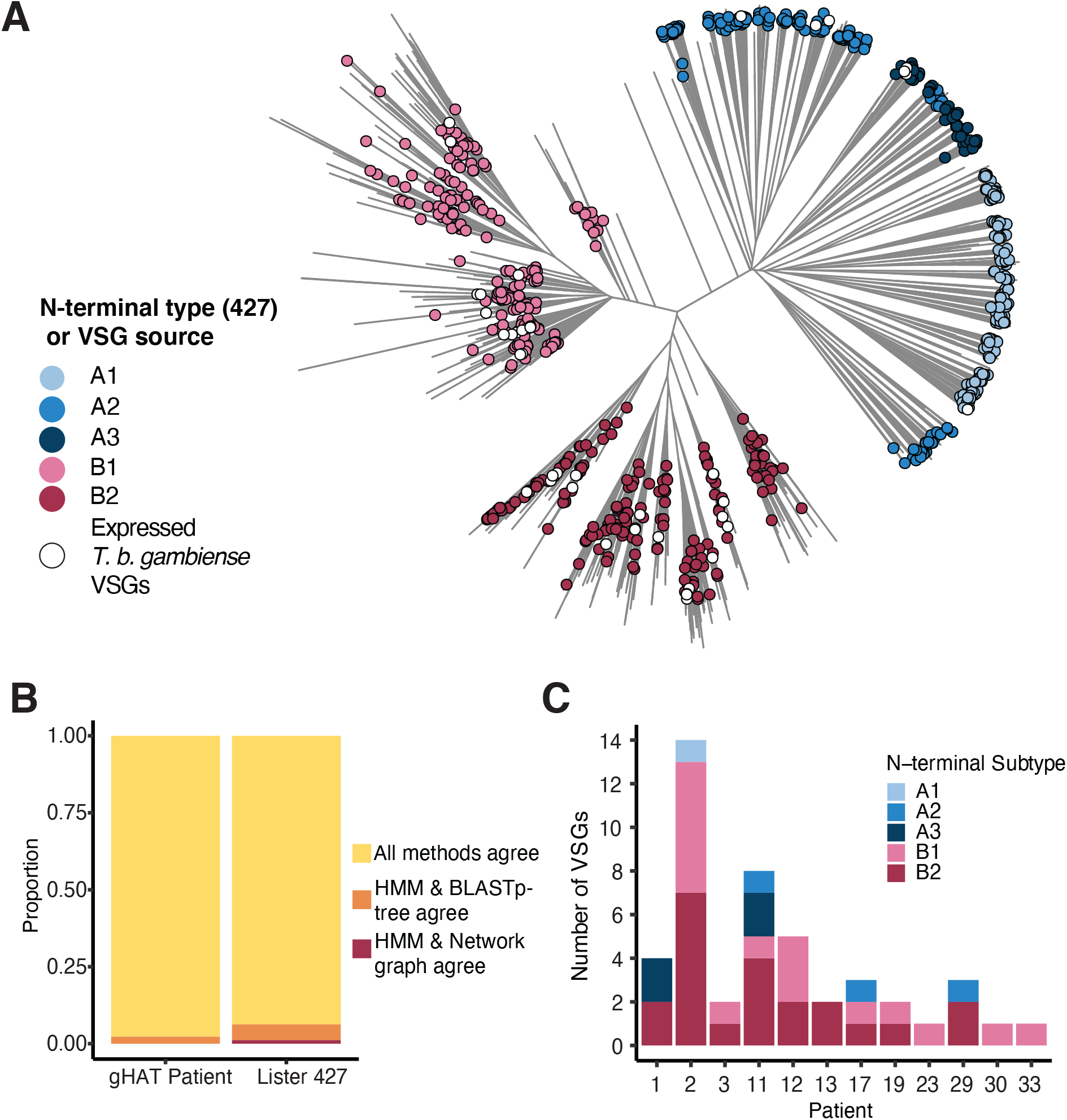
*T. b. gambiense* samples show a bias towards the expression of type B VSG. (A) Visualization of relatedness between N-terminal domain peptide sequences inferred by Neighbor-Joining based on normalized BLASTp scores. Legend indicates classification by HMM pipeline (for Lister 427 VSGs, to highlight agreement between the two methods) or by subspecies for *VSGs* expressed in patients. (B) Agreement between three VSG typing methods for Lister 427 *VSG* set and the expressed *T. b. gambiense* patient *VSG* set. (C) N-terminal domain subtype composition of expressed *T. b. gambiense VSGs* as determined by HMM analysis pipeline.

Both the new HMM profile and BLASTp network graph approaches generally recapitulated previous VSG classification based on BLASTp-tree, with all three methods agreeing 93.7% of the time (Figure 2B). The HMM pipeline method agreed with BLAST p-tree typing for all patient VSGs, while the network graph approach agreed for 43/44 VSGs (Figure 2B, Figure S3, Table S4 (15). It is not surprising that the HMM pipeline would better reflect the results of the BLASTp-tree method, as the N-terminal subtype HMM profiles were generated using VSGs classified by this method. Based on these data, we determined that the HMM method is a fast and accurate approach for determining the N-terminal domain types of unknown VSGs.

Our N-terminal domain typing pipeline identified the domain sequence and subtype for all 44 patient VSGs (Figure 2C). Of the expressed *T. b. gambiense* VSGs, 82% had type B N-terminal domains, and 50% or more of expressed *VSGs* within each patient were type B. This bias was not restricted to highly expressed *VSGs*, as 74.5% of all assembled *VSG* (813 of 1091 classifiable to an N-terminal subtype) were also type B.

Using the network graph approach, we also tentatively assigned C-terminal domain types to the *T. b. gambiense* VSGs (Figure S5). In line with previous observations, we saw no evidence of domain exclusion: a C-terminal domain of one type could be paired with any type of N-terminal domain (Figure S5E) (20). Most patient C-terminal domain types were type 2, while the remaining types were predominantly type 1, with only one type 3 C-terminus identified in the patient set. Overall, these data suggest that, like N-termini, expressed VSG C-termini are also biased towards certain C-terminal types. Together, these observations motivated further investigation into the VSG domains expressed during infection by other *T. brucei* subspecies. We focused this analysis on expressed N-terminal domains which make up most of the VSG protein, are more variable than C-terminal domains (15, 34), and are most likely to directly interface with the host immune system during infection (36).

### Type B VSG bias is unique to *T. b. gambiense* infection

To determine whether the bias towards type B VSGs was specific to *T. b. gambiense* infections, we analyzed RNA-seq data from a published study measuring gene expression in the blood and cerebrospinal fluid (CSF) of *T. b. rhodesiense* patients in Northern Uganda (37). These libraries were prepared conventionally after either rRNA-depletion for blood or poly-A selection for CSF samples. We analyzed only those samples for which at least 10% of reads mapped to the *T. brucei* genome. Raw reads from these samples were subjected to the VSG-seq analysis pipeline. Because the parasitemia of these patients was much higher than in our *T. b. gambiense* study, we adjusted our expression criteria accordingly to ≥0.01%, the published limit of detection of VSG-seq (26). Using this approach, we identified 77 unique *VSG* sequences across all blood and CSF samples (Figure 3A, Figure S6). SRA, the VSG-like protein that confers human serum resistance in *T. b. rhodesiense* (38), was detected in all patient samples.

**Figure 3.**
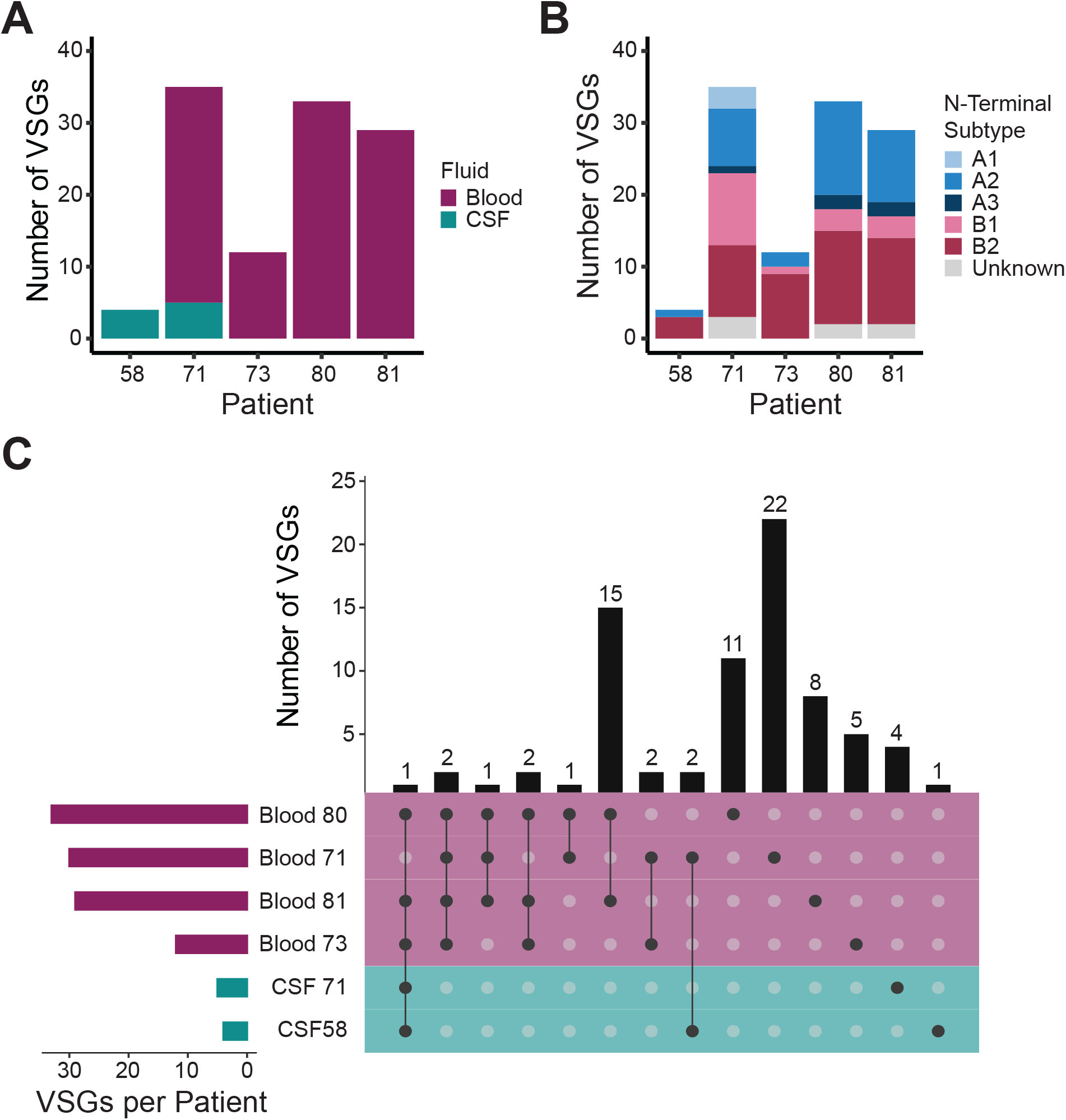
*T. b. rhodesiense* samples reveal diverse *VSG* expression but little N-terminal type bias. (A) The total number of expressed *T. b. rhodesiense VSGs* in each patient and sample type. Bar color represents the sample type from which RNA was extracted. (B) N-terminal domain subtype composition of all expressed *VSGs*. (C) Intersections of *VSGs* expressed in multiple infections. Bars on the left represent the size of the total set of VSGs expressed in each patient. Dots represent an intersection of sets, with bars above the dots representing the size of the intersection. Color indicates patient origin.

The HMM pipeline determined types for 74 of these VSG sequences; the remaining sequences appeared to be incompletely assembled, presumably due to insufficient read depth from their low level of expression. Multiple *VSGs* assembled in each patient (Figure 3A), and a large proportion of *VSGs* were expressed in multiple patients (Figure 3C). Although most VSGs detected in these patients were type B (57%, Figure 3B), this VSG type was much less predominant than in *T. b. gambiense* infection. Interestingly, *T. b. rhodesiense* patient CSF revealed another possible layer of diversity in *VSG* expression, with 5 *VSGs* expressed exclusively in this space.

### The composition of the genomic VSG repertoire is reflected in expressed VSG N-terminal domain types

One source for bias in expressed VSG type is the composition of the genomic *VSG* repertoire. To investigate the relationship between expressed *VSG* repertoires and the underlying genome composition, we took advantage of our published VSG-seq analysis of parasites isolated from mice infected with the *T. b. brucei* EATRO1125 strain. As the ‘VSGnome’ for this strain has been sequenced, we could directly compare the proportion of expressed N-terminal types to the full repertoire of types contained within the strain’s genome. In this experiment, blood was collected over time, providing data from days 6/7, 12, 14, 21, 24, 26, and 30 post-infection in all four mice, and from days 96, 99, 102, and 105 in one of the four mice (Mouse 3). Of 192 unique VSGs identified between days 0-30, the python HMM pipeline typed 190; of 97 unique VSGs identified between days 96-105, the pipeline typed 93 VSGs. The remaining VSGs were incompletely assembled by Trinity. Our analysis of VSG types over time revealed that the predominantly expressed N-terminal domain type fluctuates between type A and type B throughout the early stages of infection and in extended chronic infections (Figure S7), but the expressed *VSG* repertoire across all time points generally reflects the composition of the genomic repertoire (chi-squared p = 0.0515, Figure 4A). Parasitemia did not correlate with either the diversity of *VSG* expression or N-terminal domain type predominance (Figure S2C).

**Figure 4.**
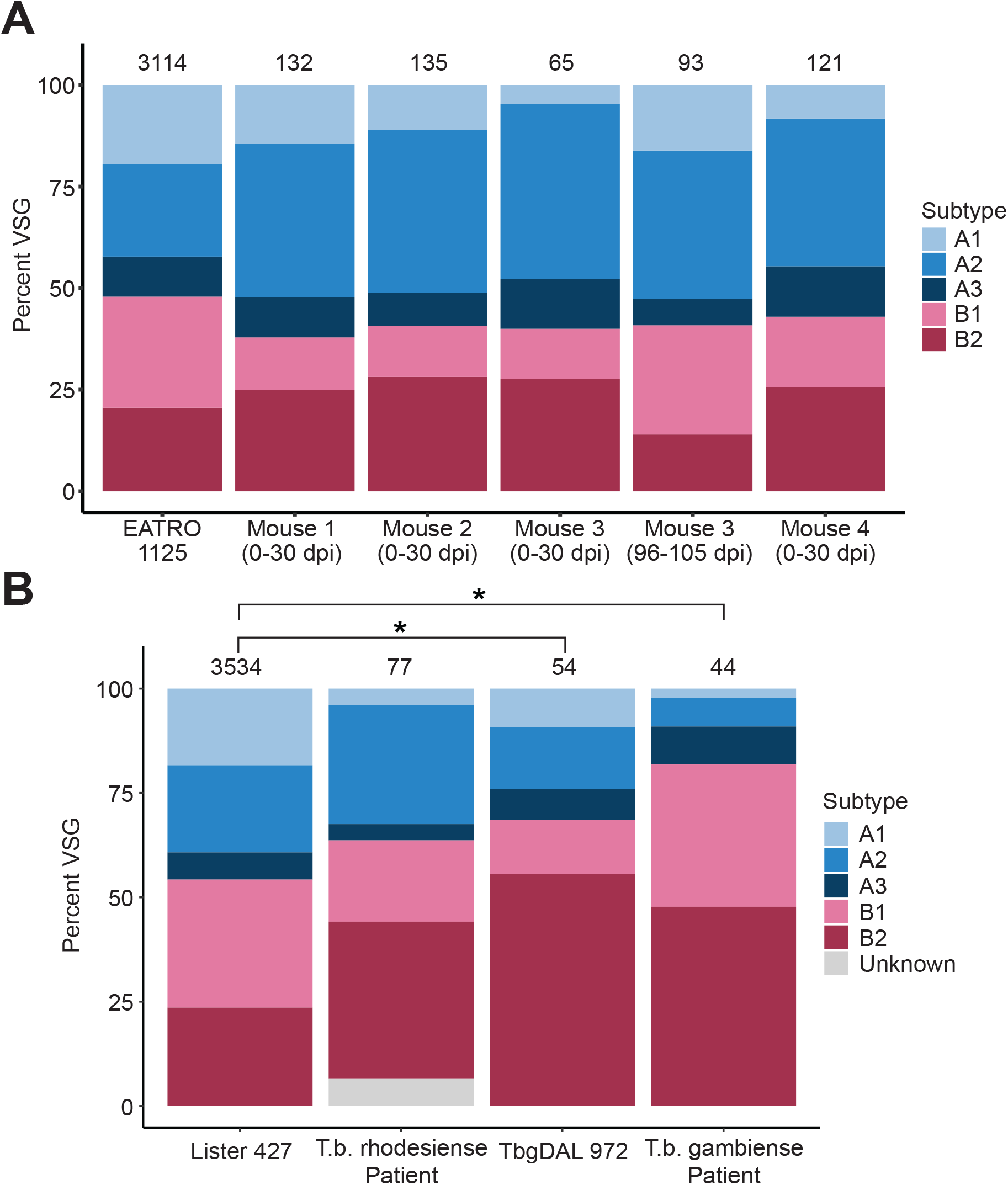
VSG expression reflects the genomic VSG repertoire of the infecting parasites. (A) Columns show the proportion of VSG types identified in each mouse infection over all time points and the proportion of VSG types in the infecting *T. b. brucei* strain, EATRO 1125. The total number of unique VSG sequences is displayed above each column. (B) A comparison of the frequencies of type A and B VSGs expressed in patients and those present in Lister 427 and DAL972 reference genomes. Relevant statistical comparisons are shown, and asterisks denote p-value < 0.05.

Unfortunately, the entire repertoire of *VSGs* encoded by most trypanosome strains is unknown, so such a direct comparison is impossible for *T. b. gambiense* and *T. b. rhodesiense* patient samples. Although the content of the ‘core’ *T. brucei* genome (containing the diploid, housekeeping genes) is similar enough among subspecies for short-read resequencing projects to be scaffolded using the TREU927 or Lister 427 reference genomes (39–41), this method cannot be applied to investigate the *VSG* repertoires of subspecies (or even individual parasite strains (27)). Because no near-complete VSGnome for any *T. b. rhodesiense* strain was available, we compared the makeup of *T. b. rhodesiense* expressed *VSGs* with the closely related and near-complete *T. b. brucei* Lister 427 repertoire (40). We observed no difference in the proportions of N-terminal types (p = 0.2422, χ^2^ test) (Figure 4B). Similarly, the proportion of N-terminal domains identified in the *T. b. gambiense* patient samples is not statistically different from the incomplete *T. b. gambiense* DAL972 genomic repertoire (p =0.0575) (Figure 4B). Both *T. b. gambiense* patient VSGs (p = 2.413e-4) and the 54 VSGs identified in *T. b. gambiense* DAL972 (p = 0.0301) have A and B type frequencies that differ significantly from the Lister427 genome. Despite limitations in the available reference genomes, together these data support a model in which VSG types are drawn from the repertoire at a roughly equal frequency to their representation in the genome, with *T. b. gambiense* exhibiting an N-terminal type composition that differs from other subspecies.

### *VSGs* expressed by *T. b. gambiense* parasites are highly diverged from those found in the whole genome sequences of other isolates

We sought to understand how the *VSGs* expressed in the *T. b. gambiense* patient isolates related to known *T. b. gambiense VSG* sequences and whether there was evidence of recombination within the expressed *VSGs*. Initial attempts to BLAST the assembled *VSGs* against the DAL972 whole genome assembly provided very few hits even using extremely permissive settings (-word_size 11 -evalue 0.1). This was unexpected but may reflect the relatively low coverage of the total VSG repertoire in the DAL972 genome assembly, which primarily covers the ‘core’ genome.

To evaluate the relationship between the expressed *VSGs* and other isolates, we took advantage of publicly available short-read whole genome datasets for 36 *T. b. gambiense* strains from three groups defined by their region and date of isolation: Côte d’Ivoire 1980’s, Côte d’Ivoire 2000’s, and DRC 2000’s (42, 43). We searched for similarity between the expressed *VSGs* and each isolate genome by mapping short reads to each assembled expressed *VSG*: regions in which reads align to a specific VSG are present somewhere in the genome of the isolate, while regions with no alignments must either be unique to gHAT patients or sufficiently diverged to no longer map.

Using representative genes from the model organisms *C. elegans, D. melanogaster*, and *E. coli* as negative controls and *T. b. gambiense* GAPDH as a positive control, we determined the appropriate read length for evaluating sequence representation. The majority of each negative control gene (66.3% average across all controls) was covered by a successful alignment using 20 bp sequences and allowing 2 or fewer mismatches (Figure S8A), indicating that read mapping at this length is not sufficiently specific. Increasing the sequence query length to 30bp greatly decreased mapping to the negative controls, such that an average of 1.4% of each gene was represented within the genomic datasets. The *T. b. gambiense* GAPDH control, on the other hand, retained 100% read coverage across the whole gene at all read lengths (Figure S8B). Thus, a 30 bp query is of appropriate stringency to measure the sequence representation of the patient *VSGs* within the whole genome datasets.

Using this query length, ~70% of the patient VSG ORF on average was absent from each genome dataset (Figure S9). Further analysis showed that C-terminal domain sequences were well represented within all genomic datasets regardless of origin (mean mapped read coverage = 77.4%), while there was relatively little nucleotide sequence similarity between the isolate genomes and the N-termini expressed by parasites in gHAT patients (16.4%, Figure 5A). Aligned nucleotide coverage was significantly higher for the genomic datasets from strains also isolated in the DRC (where the gHAT patients originated) than those isolated in Côte d’Ivoire from either time period (Figure 5B), suggesting a geographic component to *VSG* repertoires. Nonetheless, nucleotide coverage was still very low for DRC isolates when mapping to expressed N-termini (18.4%) with no expressed VSG entirely present within the genomic datasets.

**Figure 5.**
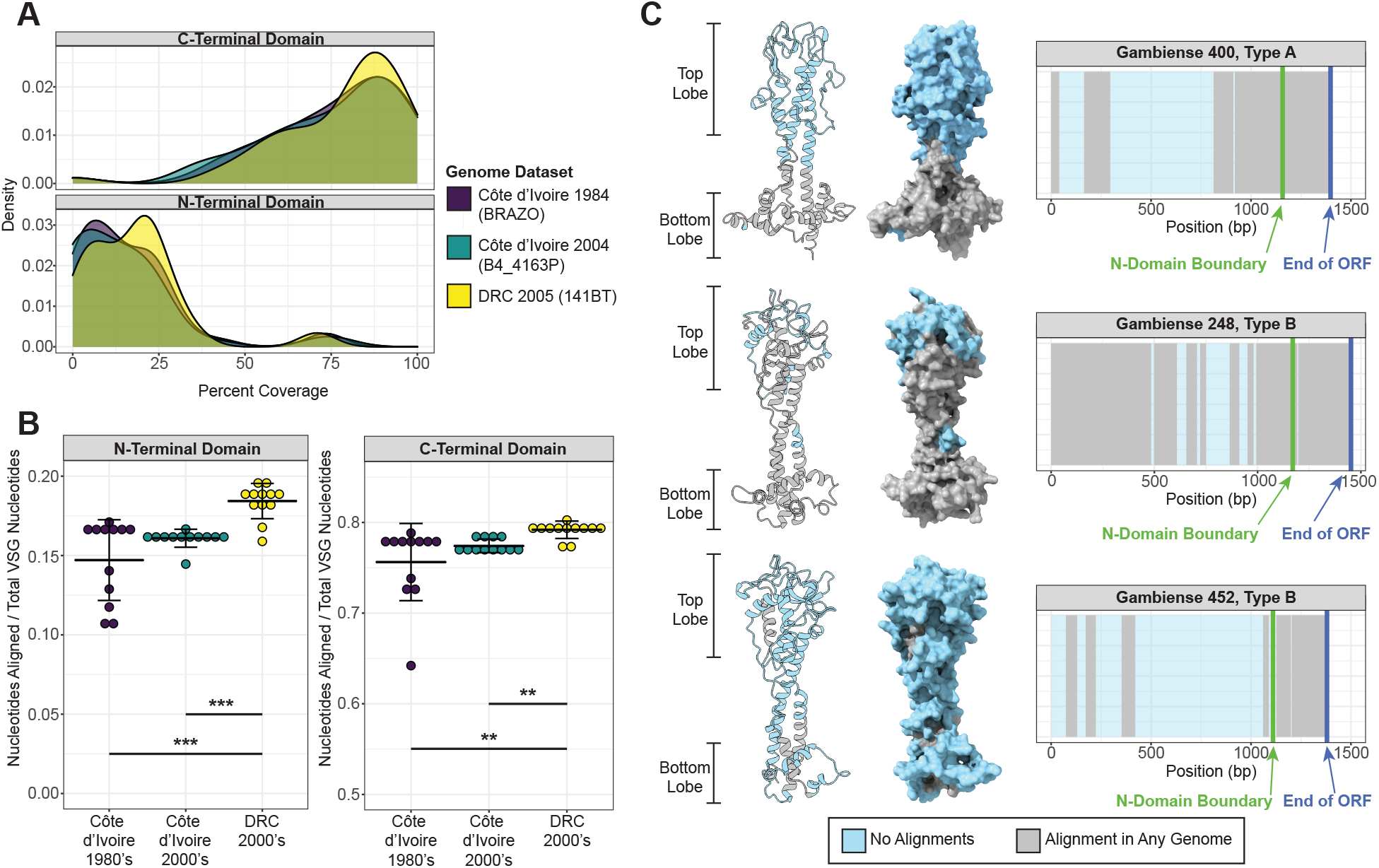
Diversification is most dramatic in exposed regions of the VSG. A) Density plot showing the percentage of each of the patient VSG ORF sequence that had at least one whole genome sequencing read (30bp length) align for each of three representative whole genome datasets (n = 12 per group). The average coverage is shown by a vertical line. B) Plots comparing sequence representation within the patient VSG N-terminal and C-terminal domains for each group. Representation for each VSG is quantified as the proportion of nucleotides in each domain with at least one alignment to the total number of nucleotides in that domain, with the average representation of all VSGs for each genome shown. Crossbars indicate mean and standard deviation within group. Significant differences between groups were determined using Kruskal-Wallis followed by a post-hoc Dunn’s test (** = p-vaue < 0.01, *** = p-value < 0.001). C) Models showing the predicted N-terminal domain structures of the three patient VSGs. The VSG shown are the type A (Gambiense 248) and type B (Gambiense 452) VSGs with highest reported ORF coverage, and a type B VSG (Gambiense 452) with average ORF coverage. Monomer structures are oriented so the polymerization interface is away from the viewer. To the right of each model is a map of coverage across each *VSG* ORF. Regions with at least one alignment from any of the 36 genomic datasets are shown in gray, and regions with no alignment are shown in blue.

To understand where diverged sequences occurred on the VSG protein, we modeled the regions of sequence divergence on predicted N-terminal domain monomer structures of each patient VSG. Strikingly, we found that the DNA sequences that encoded residues in the top lobe of the protein were invariably absent from all genomic datasets (Figure 5C). Overall, this analysis indicates that the *VSGs* expressed in the *T. b. gambiense* patient isolates are highly diverged from those within the DAL972 genome as well as from other sequenced field isolates, particularly within the parts of N-terminal domain most likely to interface with host antibody. These results are also consistent with geographic variation in *T. b. gambiense VSG* repertoires.

## Discussion

African trypanosomes evade the host adaptive immune response through a process of antigenic variation where parasites switch their expressed *VSG* (44). The genome of *T. brucei* encodes a large repertoire of *VSG* genes, pseudogenes, and gene fragments that can be expanded continuously through recombination to form entirely novel “mosaic”*VSGs* (17). While antigenic variation has been studied extensively in culture and animal infection models, our understanding of the process in natural infections, particularly human infection, is limited. Most experimental mouse infections are sustained for weeks to months, while humans and large mammals may be infected for several months or even years. Additionally, laboratory studies of antigenic variation almost exclusively use *T. b. brucei*, a subspecies of *T. brucei* that, by definition, does not infect humans. The primary hurdle to exploring antigenic variation in nature has been technical: it is difficult to obtain sufficient parasite material for analysis. This is especially true for infection with *T. b. gambiense*, which often exhibits extremely low parasitemia. Here we have demonstrated the feasibility of VSG-seq to analyze *VSG* expression in RNA samples isolated directly from HAT patients. Our analyses reveal unique aspects of antigenic variation in *T. b. gambiense* that can only be explored by studying natural infections.

We have identified an intriguing bias towards the expression of type B VSGs in *T. b. gambiense* infection, which appears to be specific to this *T. brucei* subspecies. Comparison of expressed *VSG* repertoires to publicly available genomic *VSG* repertoires suggests that the genomic *VSG* repertoire determines the distribution of VSG N-terminal types expressed during *T. brucei* infection. Thus, the *T. b. gambiense VSG* repertoire may contain a larger proportion of type B VSGs than its more virulent counterparts. Could a bias towards certain VSG types, whether due to a difference in repertoire composition or expression preference, account for unique features of *T. b. gambiense* infection, including its chronicity and primarily anthroponotic nature (45)?

Little is known about how differences in VSG proteins relate to parasite biology or whether there could be biological consequences to the expression of specific VSG N- or C-terminal types. Type A *var* genes in *Plasmodium falciparum* infection are associated with severe malaria (46–50), and similar mechanisms have been hypothesized to exist in *T. vivax* and *T. congolense* infections (51–54). In *T. brucei*, several VSGs have evolved specific functions besides antigenic variation (54). The first type B VSG structure was recently solved (55), revealing a unique *O*-linked carbohydrate in the VSG’s N-terminal domain that interfered with the generation of protective immunity in a mouse infection model. Perhaps structural differences between each VSG type, including glycosylation patterns, could influence infection outcomes. Further research will be needed to determine whether the observed predominance of type B VSGs could influence the biology of *T. b. gambiense* infection.

Another possibility we cannot rule out, however, is that the gHAT samples are biased due to selection by the serological test used for diagnosis. Patients were screened for *T. b. gambiense* infection using the CATT, a serological test that uses parasites expressing VSG LiTat 1.3 as an antigen. LiTat 1.3 contains a type B2 N-terminal domain (56, 57). Patients infected with parasites predominantly expressing type B VSGs may be more likely to generate antibodies that cross-react with LiTat1.3, resulting in preferential detection of these cases. In contrast, *T. b. rhodesiense* can only be diagnosed microscopically, removing the potential to introduce bias through screening. It remains to be investigated whether samples from patients diagnosed using newer screening tests, which include the invariant surface glycoprotein ISG65 and the type A VSG LiTat 1.5 (23), would show similar bias towards the expression of type B VSGs.

Such a bias, if it exists, would be important to understand, as it could affect the ability to detect a subset of gHAT infections. The diversity and corresponding divergence of expressed VSGs from publicly available genomic sequences could have similar implications. Although diversity in *T. b. gambiense* infection appeared lower overall than previous measurements from experimental mouse infections (17, 18, 26), the correlation we observed between parasitemia and diversity in *T. b. gambiense* isolates suggests that our sampling was incomplete. Indeed, in our analysis of *T. b. rhodesiense* infection (a more reasonable comparison to mouse infection given similar expression cutoffs and parasitemia), we observed diversity similar to or higher than what has been observed in *T. b. brucei* mouse infections. Moreover, *T. b. rhodesiense* patient CSF revealed another layer of diversity in *VSG* expression, with 5 *VSGs* expressed exclusively in this space. Although this observation is difficult to interpret without information about the precise timing of sample collection, a recent study in mice showed that extravascular spaces harbor much of the antigenic diversity during infection (58). It is exciting to speculate that different organs or body compartments could harbor different sets of VSGs in humans as well.

Overall, our analysis of *VSG* expression in *T. b. gambiense* and *T. b. rhodesiense* patients confirmed the long-held assumption that *VSG* diversity is a feature of natural infection. One potential consequence of this striking diversity is that the genomic *VSG* repertoire might be exploited very rapidly, creating pressure for the parasite to diversify its *VSG* repertoire as the mammalian host generates antibodies against each expressed *VSG*. Our results are consistent with this, revealing extreme divergence in the patient *VSGs* from 36 publicly available *T. b. gambiense* whole genome sequencing datasets. Even when mapping relatively short 30bp genomic sequences to each VSG, we could only find evidence for ~30% of each *VSG* ORF. Without assembled genomes, it is difficult to infer recombination patterns or mechanisms from this analysis. The fact that only very short stretches of homology could be found within the N-terminal domain, however, is consistent with recombination through microhomology-mediated end joining, a DNA repair mechanism that uses short stretches of homology (5-20bp) to repair DNA damage (59). This appears to be the favored form of DNA repair in the *VSG* expression site and has been hypothesized to play a role in *VSG* switching (59, 60). The data presented here suggest this mechanism, or a similar one, may play a role in diversification of the *VSG* repertoire as well.

We also observed divergence between geographically separate parasite populations. Past research has shown that the sensitivity of serological tests for gHAT, which detect antibodies against the LiTat 1.3 VSG, vary regionally, potentially due to differences in the underlying genomic or expressed *VSG* repertoire in circulating strains (56, 57). Our data is consistent with such a possibility, with the *VSGs* expressed in patients from the DRC sharing more sequence similarity with isolates from the same country than those from Côte d’Ivoire. Geographic variation has been observed in *var* gene repertoires of *Plasmodium falciparum* (61) and the *VSG* repertoire of *Trypanosoma vivax*, another African trypanosome (53). A better understanding of such differences in *T. brucei* could inform the development of future HAT diagnostics.

The positions of divergent regions within the VSG protein demonstrate the enormous pressure exerted by host antibody on the repertoire of *T. b. gambiense*. While the C-termini of patient *VSGs* were well-represented, the majority of each N-terminal sequence was undetectable in the 36 genomes we analyzed. Notably, in even the most conserved VSG N-termini, sequences encoding the top lobe of the VSG were completely absent from the genomes we analyzed. VSG proteins are packed together very closely on the parasite cell surface, presumably preventing host antibody from accessing epitopes close to or within the C-terminus (36). Thus, those regions with no nucleotide similarity correspond directly to the parts of the VSG protein most likely to be exposed to host antibody.

In addition to confirming that certain aspects of antigenic variation observed in experimental *T. brucei* infection are features of natural infection, this study has revealed unique features of the process in *T. b. gambiense*. This subspecies appears to preferentially express certain VSG N-termini, which could be related to the unique biology of the parasite. Additionally, wild *VSG* repertoires may be more diverse than previously expected with potential geographic variation. While mouse models can recapitulate certain aspects of the process, new biology remains to be uncovered by studying antigenic variation in its natural context.

## Methods

### Ethics statement

The blood specimens from *T.b. gambiense* infected patients were collected within the projects, “Longitudinal follow-up of CATT seropositive, trypanosome negative individuals (SeroSui)” and “An integrated approach for identification of genetic determinants for susceptibility for trypanosomiasis (TrypanoGEN)” (62). In France, the SeroSui study received approval from the Comité Consultatif de Déontologie et d’Ethique (CCDE) of the French National Institute for Sustainable Development Research (IRD), May 2013 session. In Belgium, the study received approval from the Institutional Review Board of the Institute of Tropical Medicine (reference 886/13) and the Ethics Committee of the University of Antwerp (B300201318039). In the Democratic Republic of the Congo, the projects SeroSui and TrypanoGEN were approved by the Ministry of Health through the Ngaliema Clinic of Kinshasa (references 422/2013 and 424/2013). Participants gave their written informed consent to participate in the projects. For minors, additional written consent was obtained from their legal representative.

### Patient enrollment and origin map

Patients originated from the DRC and were identified over six months in the second half of 2013. This identification occurred either during passive screening at the center for HAT diagnosis and treatment at the hospital of Masi Manimba, or during active screening by the mobile team of the national sleeping sickness control program (PNLTHA) in Masi Manimba and Mosango health zones (Kwilu province, DRC).

Individuals were screened for the presence of specific antibodies in whole blood with the CATT test. For those reacting blood positive in CATT, we also tested twofold serial plasma dilutions of 1/2-1/32 were also tested and determined the CATT end titer was determined. CATT positives underwent parasitological confirmation by direct microscopic examination of lymph (if enlarged lymph nodes were present), and examination of blood by the mini-anion exchange centrifugation technique on buffy coat (63). Individuals in whom trypanosomes were observed underwent lumbar puncture. The cerebrospinal fluid was examined for white blood cell count and the presence of trypanosomes to determine the disease stage and select the appropriate treatment. Patients were questioned about their place of residence. The geographic coordinates of their corresponding villages were obtained from the Atlas of HAT (64) and plotted on a map of the DRC using ArcGIS^®^ software by Esri. Distances were determined and a distance matrix generated (see Supplemental Table 2).

### Patient blood sample collection and total RNA isolation

A 2.5 mL volume of blood was collected from each patient in a PAXgene Blood RNA Tube. The blood was mixed with the buffer in the tube, aliquoted in 2 mL volumes and frozen in liquid nitrogen for a maximum of two weeks. After arrival in Kinshasha, tubes were stored at −70°C. Total RNA was extracted and isolated from each blood sample as previously described (65).

### Estimation of parasitemia

Two approaches were used to estimate parasitemia. First, a 9 mL volume of blood on heparin was centrifuged, 500 microliters of the buffy coat were taken up and trypanosomes were isolated using the mini-anion exchange centrifugation technique. After centrifugation of the column eluate, the number of parasites visible in the tip of the collection tube were estimated. Second, Spliced Leader (SL) RNA expression levels were measured by real-time PCR as previously described (65). A Ct value was determined for each patient blood sample. Real-time PCR was performed on RNA samples before reverse transcription to verify the absence of DNA contamination.

### RNA sequencing

DNase I-treated RNA samples were cleaned up with 1.8× Mag-Bind TotalPure NGS Beads (Omega Bio-Tek, # M1378-01). cDNA was generated using the SuperScript III First-strand synthesis system (Invitrogen, 18080051) according to manufacturer’s instructions. 8 microliters of each sample (between 36 and 944 ng) were used for cDNA synthesis, which was performed using the oligo-dT primer provided with the kit. This material was cleaned up with 1.8x Mag-Bind beads and used to generate three replicate library preparations for each sample. These technical replicates were generated to ensure that any *VSGs* detected were not the result of PCR artifacts(66, 67).

Because we expected a low number of parasites in each sample, we used a nested PCR approach to prepare the VSG-seq libraries. First, we amplified *T. brucei* cDNA from the parasite/host cDNA pool by PCR using a spliced leader primer paired with an anchored oligo-dT primer (SL-1-nested and anchored oligo-dT; Supplemental Table 1). 20 cycles of PCR were completed (55°C annealing, 45s extension) using Phusion polymerase (Thermo Scientific, #F530L). PCR reactions were cleaned up with 1.8× Mag-Bind beads. After amplifying *T. brucei* cDNA, a *VSG*-specific PCR reaction was carried out using M13RSL and 14-mer-SP6 primers (see primers; Supplemental Table 1). 30 cycles of PCR (42°C annealing, 45s extension) were performed using Phusion polymerase. Amplified *VSG* cDNA was then cleaned up with 1X Mag-Bind beads and quantified using a Qubit dsDNA HS Assay (Invitrogen Q32854).

Sequencing libraries were prepared from 1 ng of each VSG PCR product using the Nextera XT DNA Library Preparation Kit (Illumina, FC-131-1096) following the manufacturer’s protocol except for the final cleanup step, which was performed using 1X Mag-Bind beads. Single-end 100bp sequencing was performed on an Illumina HiSeq 2500. Raw data are available in the National Center for Biotechnology Information (NCBI) Sequence Read Archive under accession number PRJNA751607.

### VSG-seq analysis of *T. b. gambiense* and *T. b. rhodesiense* sequencing libraries

To analyze both *T. b. gambiense* (VSG-seq preparations) and *T. b. rhodesiense* (traditional mRNA sequencing library preparations; sequences were obtained from ENA, accession numbers PRJEB27207 and PRJEB18523), we processed raw reads using the VSG-seq pipeline available at https://github.com/mugnierlab/VSGSeqPipeline. Briefly, *VSG* transcripts were assembled *de novo* from quality- and adapter-trimmed reads for each sample (patient or patient replicate) from raw reads using Trinity (version 5.26.2) (68). Contigs containing open reading frames (ORFs) were identified as previously described (26). ORF-containing contigs were compared to Lister 427 and EATRO1125 *VSGs* as well as a collection of known contaminating non-*VSG* sequences. Alignments to *VSGs* with an E-value below 1×10^−10^ that did not match any known non-*VSG* contaminants were identified as *VSG* transcripts. For *T. b. gambiense* replicate libraries, *VSG* ORFs identified in any patient replicate were consolidated into a sole reference genome for each patient using CD-HIT (version 4.8.1) (69) with the following arguments: -d 0 -c 0.98 -n 8 -G 1 -g 1 -s 0.0 -aL 0.0. Final *VSG* ORF files were manually inspected.

Two *T. b. gambiense* patient *VSG*s (Patients 11 and 13) showed likely assembly errors. In one case, a *VSG* was duplicated and concatenated, and in another, two *VSGs* were concatenated.

These reference files were manually corrected (removing the duplicate or editing annotation to reflect two *VSGs* in the concatenated ORF) so that each *VSG* could be properly quantified. *VSG* reference databases for each patient are available at https://github.com/mugnierlab/Tbgambiense2021/. For *T. b. gambiense*, we then aligned reads from each patient replicate to that patient’s consolidated reference genome using Bowtie with the parameters -*v 2 -m 1 -S* (version 1.2.3) (70).

For *T. b. rhodesiense*, we aligned each patient’s data to its own *VSG* ORF assembly. RPKM values for each *VSG* in each sample were generated using MULTo (version 1.0) (71), and the percentage of parasites in each population expressing a *VSG* was calculated as described previously (26). For *T. b. gambiense* samples, we included only *VSGs* with an expression measurement above 1% in two or more patient replicates in our analysis. For *T. b. rhodesiense* samples, we included only *VSGs* with expression >0.01%. To compare *VSG* expression between patients, despite the different reference genomes used for each patient, we used CD-HIT to cluster *VSG* sequences with greater than 98% similarity among patients, using the same parameters used to consolidate reference *VSG* databases before alignment. We gave each unique *VSG* cluster a numerical ID (e.g., Gambiense #) and chose the longest sequence within each group to represent the cluster. Before analysis, we manually removed clusters representing TgsGP and SRA from the expressed *VSG* sets. UpSet plots were made with the UpSetR package (72). The R code used to analyze resulting data and generate figures is available at https://github.com/mugnierlab/Tbgambiense2021/.

### Analysis of VSG N-terminal domains

#### Genomic *VSG* sequences

The *VSG* repertoires of *T. b. brucei* Lister 427 (“Lister427_2018” assembly), *T. b. brucei* TREU927/4 and *T. b. gambiense* DAL972 were taken from TriTrypDB (v50), while the *T. b. brucei* EATRO 1125 VSGnome was used for analysis of the EATRO1125 *VSG* repertoire (vsgs_tb1125_nodups_atleast250aas_pro.txt, available at https://tryps.rockefeller.edu/Sequences.html or GenBank accession KX698609.1 - KX701858.1). VSG sequences from other strains (except those generated by VSG-seq) were taken from the analysis in Cross, et al. (15). Likely VSG N-termini were identified as predicted proteins with significant similarity (e-value ≤ 10^−5^) to hidden Markov models (HMMs) of aligned type A and B VSG N-termini taken from (15).

#### N-terminal domain phylogenies

Phylogenies of VSG N-termini based on unaligned sequence similarities were constructed using the method described in (73) and used previously to classify VSG sequence (15). We extracted predicted N-terminal domain protein sequences by using the largest bounding envelope coordinates of a match to either type A or type B HMM. A matrix of similarities between all sequences was constructed from normalized transformed BLASTp scores as in Wickstead, et al. (73) and used to infer a neighbor-joining tree using QuickTree v1.1 (74). Trees were annotated and visualized in R with the package APE v5.2 (75).

#### HMM

For N-terminal typing by HMM, we used a python analysis pipeline available at (https://github.com/mugnierlab/find_VSG_Ndomains). The pipeline first identifies the boundaries of the VSG N-terminal domain using the type A and type B HMM profiles generated by Cross *et al*. which includes 735 previously-typed VSG N-terminal domain sequences (15). N-terminal domains are defined by the largest envelope domain coordinate that meets e-value threshold (1×10^−5^, --domE 0.00001). In cases where no N-terminal domain is identified using these profiles, the pipeline executes a “rescue” domain search in which the VSG is searched against a ‘pan-VSG’ N-terminus profile we generated using 763 previously-typed VSG N-terminal domain sequences. This set of VSGs includes several *T. brucei* strains and/or subspecies: Tb427 (559), TREU927 (138), *T. b. gambiense* DAL972 (28), EATRO795 (8), EATRO110 (5), *T. equiperdum* (4), and *T. evansi* (21). The N-terminal domain type of these VSGs were previously determined by Cross et. al (2014) by building neighbor-joining trees using local alignment scores from all-versus-all BLASTp similarity searches (15). Domain boundaries are called using the same parameters as with the type A and B profiles.

After identifying boundaries, the pipeline extracts the sequence of the N-terminal domain, and this is searched against five subtype HMM profiles. To generate N-terminal domain subtype HMM profiles, five multiple sequence alignments were performed using Clustal Omega (76) with the 763 previously-typed VSG N-terminal domain sequences described above; each alignment included the VSG N-terminal domains of the same subtype (A1, A2, A3, B1, and B2). Alignment output files in STOCKHOLM format were used to generate distinct HMM profiles for type A1, A2, A3, B1, and B2 VSGs using the pre-determined subtype classifications of the 763 VSGs using HMMer version 3.1b2 (77). The number of sequences used to create each subtype profile ranged from 75 to 211. The most probable subtype is determined by the pipeline based on the highest scoring sequence alignment against the subtype HMM profile database when HMMscan is run under default alignment parameters. The pipeline generates a FASTA file containing the amino acid sequence of each VSG N-terminus and a CSV with descriptions of the N-terminal domain including its type and subtype.

#### Network graph

N-terminal network graphs were made using VSG N-terminal domains from the TriTrypDB Lister427_2018 and *T. b. gambiense* DAL972 (v50) VSG sets described above, and the *T. b. gambiense* and *T. b. rhodesiense* patient VSG N-termini which met our expression thresholds. Identified N-terminal domains were then subjected to an all-versus-all BLASTp. A pairwise table was created that includes each query-subject pair, the corresponding alignment E-value, and N-terminal domain type of the query sequence if previously typed in Cross, et al. (15). Pseudogenes and fragments were excluded from the Lister427_2018 reference prior to plotting by filtering for *VSG* genes annotated as pseudogenes and any less than 400 amino acids in length, as the remaining sequences are most likely to be full length VSG. Network graphs were generated with the igraph R package(78) using undirected and unweighted clustering of nodes after applying link cutoffs based on E-value 10^−2^. The leading eigenvector clustering method (35) was used to detect and assign nodes to communities based on clustering (cluster_leading_eigen() method in igraph).

### Analysis of VSG C-terminal domains

VSG C-termini were extracted from expressed *T. b. gambiense VSGs*, *T.b. gambiense* DAL972 (v50), and 545 previously-typed VSG C-termini from the Lister 427 strain using the C-terminal HMM profile generated by Cross et al. (15) and the same HMMscan parameters as for N-termini (E-value < 1×10^−5^; largest domain based on envelope coordinates). An all-vs-all BLASTp was performed on these sequences, and network graphs were generated in the same manner as the N-terminal network graphs. Links were drawn between C-termini with a BLASTp E-value 1×10^−3^. The leading eigenvector method for clustering (35) was used to detect and assign nodes to communities based on clustering (cluster_leading_eigen() method in igraph).

### Comparison of gHAT patient VSGs to sequenced whole genomes of *T.b. gambiense* isolates

Publicly available whole genome Illumina sequencing reads for 24 *T.b. gambiense* isolates from Côte d’Ivoire were fetched from the ENA database and 12 datasets for isolates from the DRC were downloaded from DataDryad. All datasets analyzed exist as raw sequencing reads and do not have associated ORF assemblies or VSG gene annotations. We therefore determined the presence or absence of sequences similar to patient VSG by alignment. Raw reads were adapter and quality trimmed using Trim_Galore (version 0.5.0) under default parameters and truncated to desired query lengths of 20, 30, and 50 bp using Trimmomatic (79) (version 0.38) ‘CROP’ option. Whole genome sequence datasets were aligned to the assembled patient VSG nucleotide sequences using Bowtie with the parameters -*v 2 -a -S* (version 1.1.1). Bowtie does not support gapped alignments and the number of mismatched bases per read can be adjusted to control the stringency of alignments, therefore this aligner was used to assess the size of regions of sequence similarity between the patient VSG and genomic sequences. Bedtools (80) (version 2.27.0) genomecov was used to summarize alignment coordinates and read depth for downstream analysis. Alignment ranges were plotted with the IRanges R package(81). Patient VSG gene coverage was calculated as the regions of sequence with an aligned read depth of at least one divided by the full ORF sequence or domain length in bp.

To model regions of sequence divergence and similarity, the secondary structures for each of the 44 gHAT patient VSG were predicted using Phyre2 (82) batch processing under default parameters. Automated threading returned hits to VSG N-terminal domain chain templates from the PDB with 100% confidence for all patient VSG. Predicted structures were visualized and figures generated in ChimeraX (83).

## Supporting information

Supplemental Table 1. Primer sequences.

Supplemental Table 2. gHAT patient distance matrix.

Supplemental Table 3. gHAT VSG expression data.

Supplemental Table 4. Tables comparing BLAST-tree, HMMscan, and network plot typing methods.

Supplemental Table 5. rHAT VSG expression data.

Supplemental Figure 4. BLASTp-tree of all T. b. gambiense VSGs. File attached.

## Acknowledgments

We are very grateful to the patients without whom this work would not have been possible. We thank George Cross and Danae Schulz for comments on the manuscript, and Mary Gebhardt for help with GIS. The Atlas of HAT is an initiative of the World Health Organization (WHO), jointly implemented with the Food and Agriculture Organization of the United Nations (FAO) in the framework of the Programme Against African Trypanosomiasis (PAAT). Field work and specimen collection in DRC were funded through the Wellcome Trust (study number 099310/Z/12/Z) awarded to the TrypanoGEN Consortium (www.trypanogen.net), members of H3Africa (h3africa.org). Sample work-up was supported by the Research Foundation Flanders (FWO grant 1501413N). Work by BW was supported by University of Nottingham/Wellcome Trust Institutional Strategic Support Fund award 204843/Z/16/Z. MRM and SS were supported by Office of the Director, NIH (DP5OD023065). JS is supported by NIH T32AI007417.

## Supplement

Supplemental Table 1. Primer sequences.

Supplemental Table 2. gHAT patient distance matrix.

Supplemental Table 3. gHAT VSG expression data.

Supplemental Table 4. Tables comparing BLAST-tree, HMMscan, and network plot typing methods.

Supplemental Table 5. rHAT VSG expression data.

**Supplemental Figure 1.**
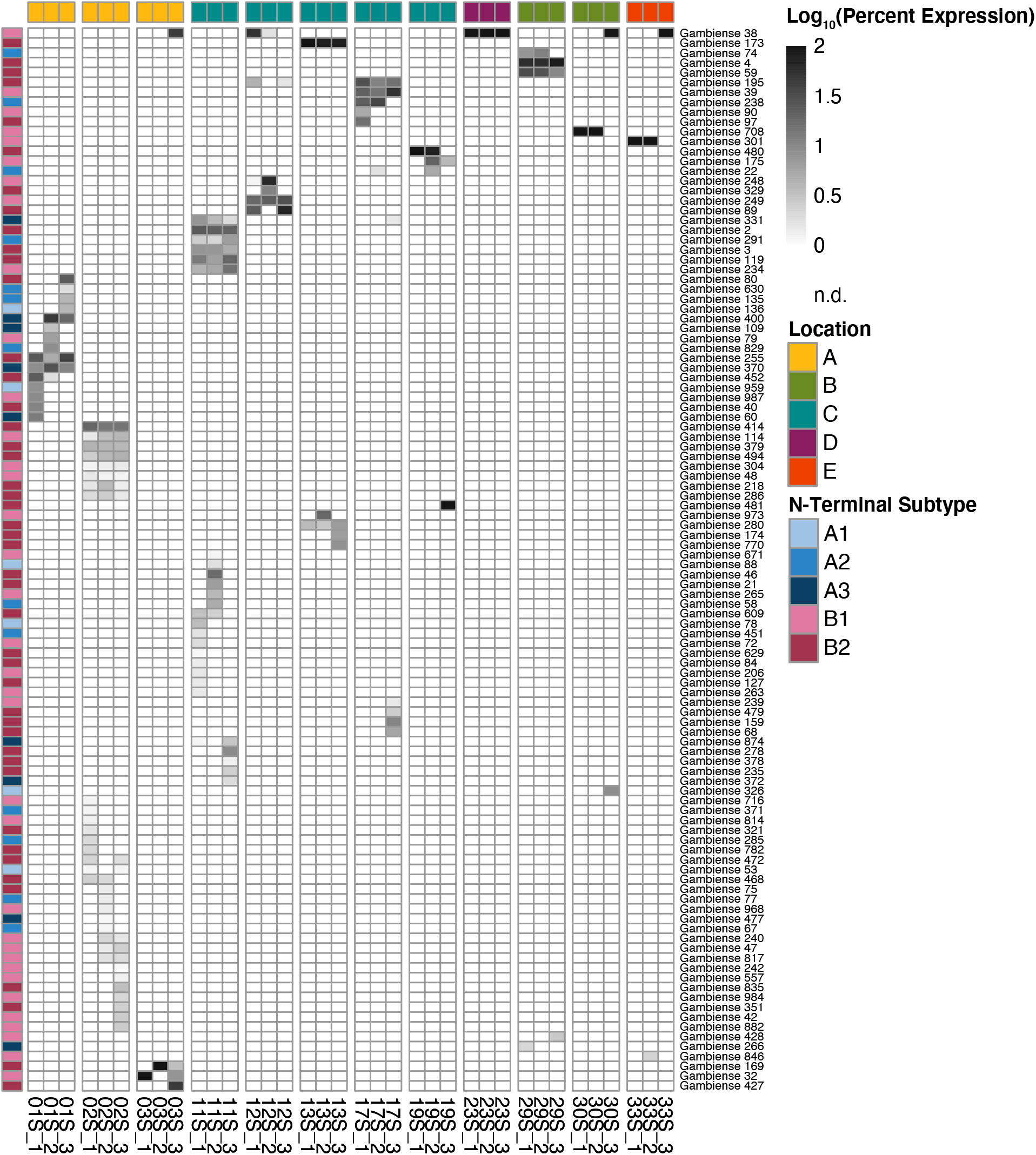
Heatmap of all assembled *T.b. gambiense* patient VSGs. Greyscale shows log_10_ of the estimated percentage of the parasite population expressing each VSG. Variants expressed by less than 1% of parasites considered not detected (n.d.).

**Supplemental Figure 2.**
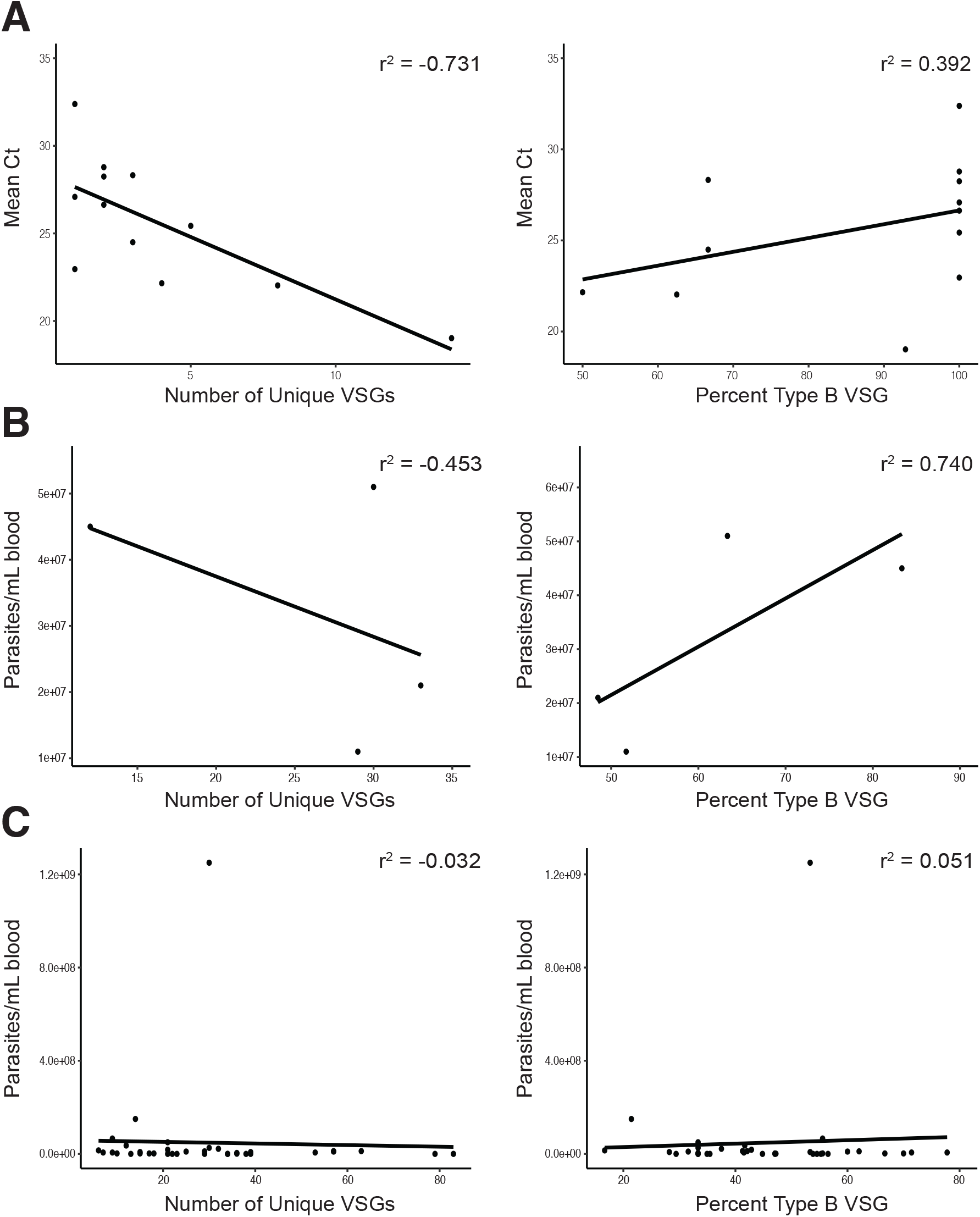
Correlation between parasitemia and diversity and N-terminal type distribution. (A) Correlation plots for *T.b. gambiense* infected patients. (B) Correlation plots for *T.b. rhodesiense* infected patients from Mulindwa et al. 2018. (C) Correlation plots for VSG diversity and percent of N-terminal domain type B for *T.b. brucei* infected mice from Mugnier et al. 2015.

**Supplemental Figure 3.**
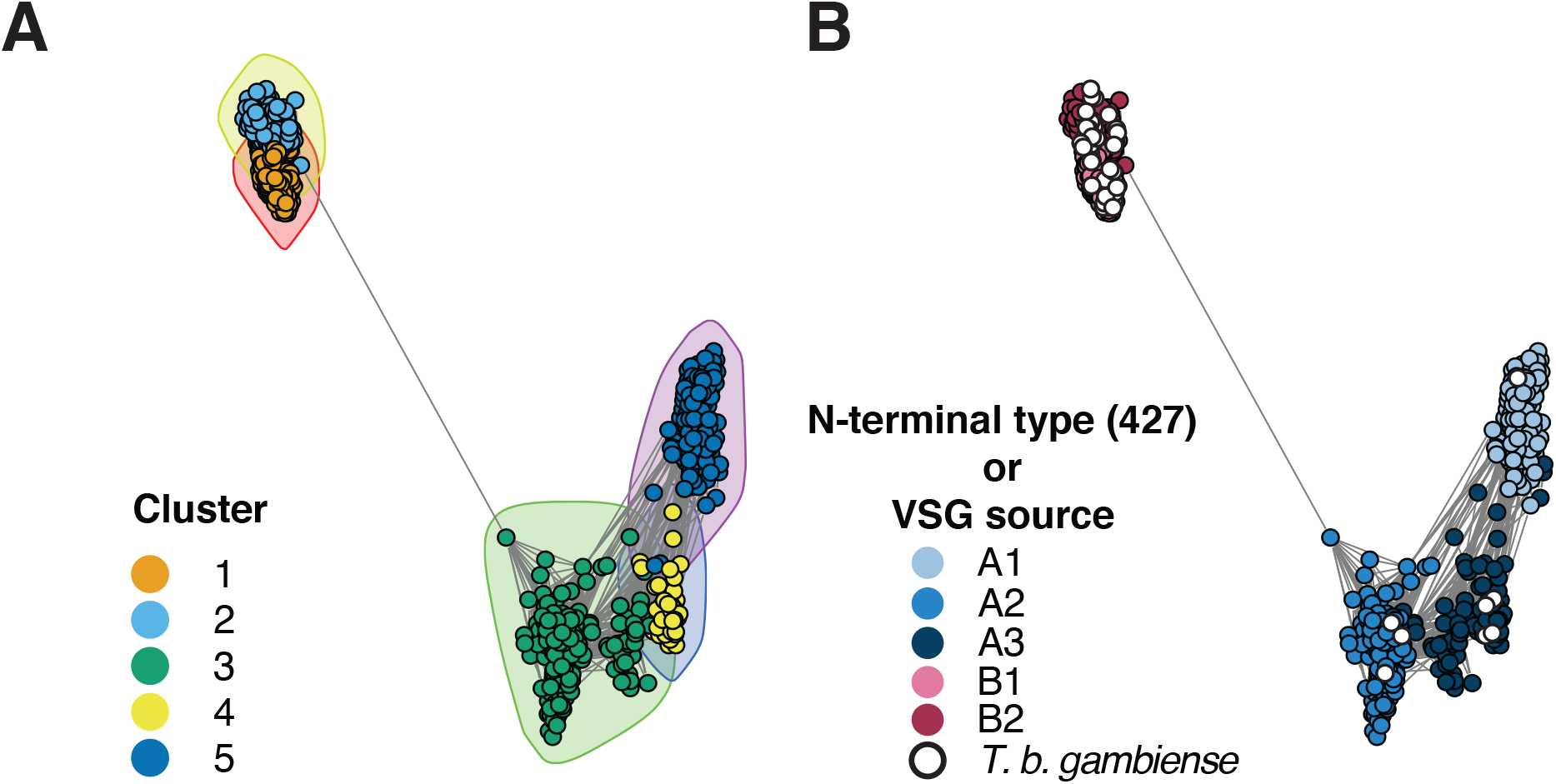
(A) Network plot showing peptide sequence relatedness between N-terminal domains. Each point represents a VSG N-terminus. A link was drawn between points if the BLASTp e-value was less than 10^−2^. Colors and shaded circles represent community assignments determined by the clustering algorithm. (B) The same graph as in (A), but points are manually colored by known N-terminal subtype from Cross et al. or by subspecies for VSGs identified in patients.

**Supplemental Figure 4. BLASTp-tree of all *T. b. gambiense VSGs*.** File attached.

**Supplemental Figure 5.**
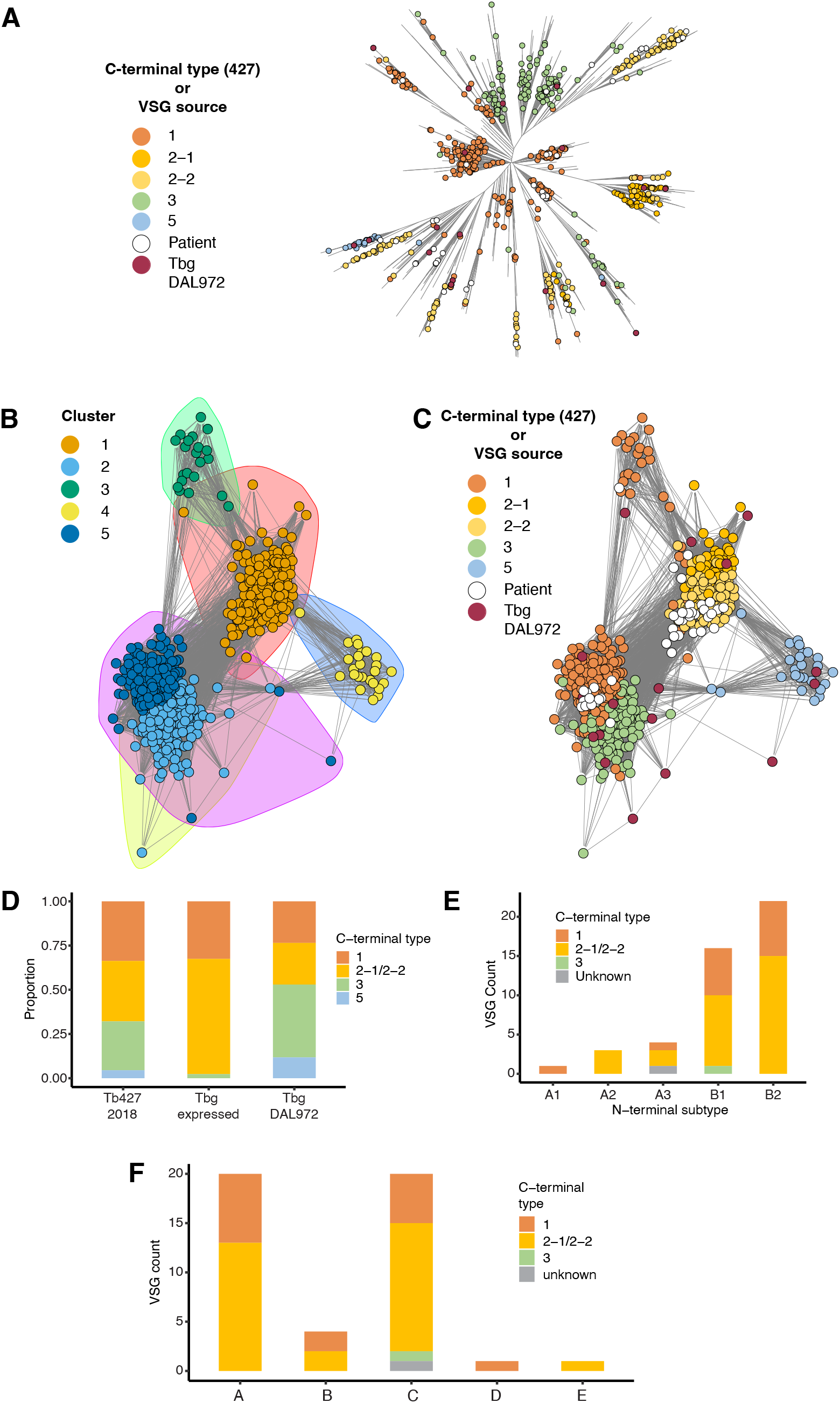
Expressed VSG C-termini are primarily type 1 and type 2. A) BLASTp-tree of C-terminal domains. Points are colored based on previously determined C-terminal type from Cross et al. or by the source of the sequence (genomic or expressed) for *T. b. gambiense VSGs*. B) Network plot showing peptide sequence relatedness between C-terminal domains in *T. b. gambiense* expressed VSGs. Each point represents a VSG C-terminus. A link was drawn between points if the BLASTp e-value was less than 1×10^3^. Points are colored by the cluster determined by the clustering algorithm. Shaded circles also indicate clusters. C) Same network plot as in B but colored by previously determined C-terminal type from Cross et al., or by source for unclassified genomic or expressed *VSGs*. D) VSG C-terminal types, based on cluster assignment visualized in panel B, in genomic and expressed *VSG* sets. E) Pairing of C- and N-termini in *T. b. gambiense* patients. F) C-termini detected in each patient village.

**Supplemental Figure 6.**
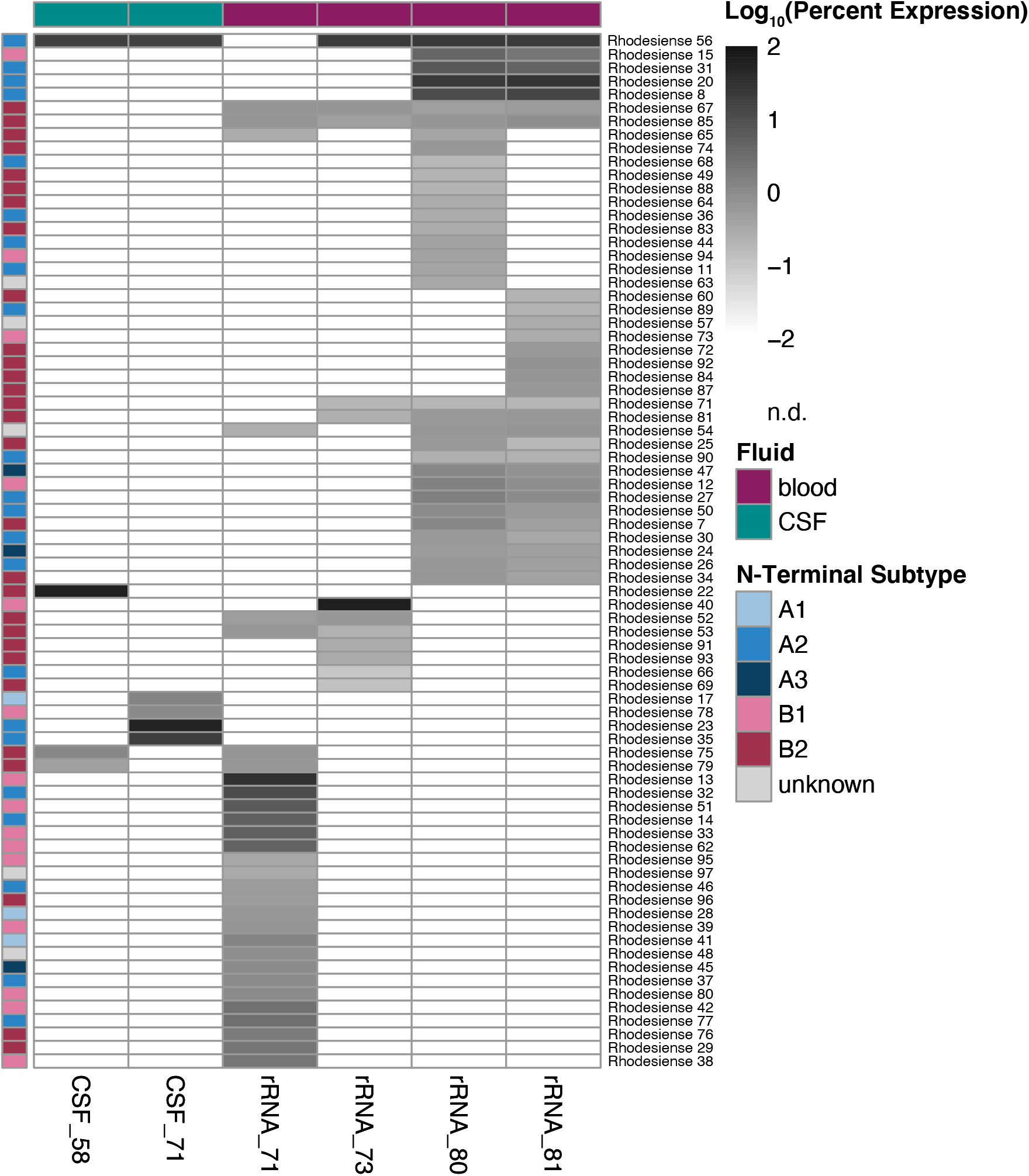
Heatmap of all assembled *T. b. rhodesiense* patient VSGs. Greyscale shows log_10_ of the estimated percentage of the parasite population expressing each VSG. Variants expressed by less than 0.01% of parasites considered not detected (n.d.).

**Supplemental Figure 7.**
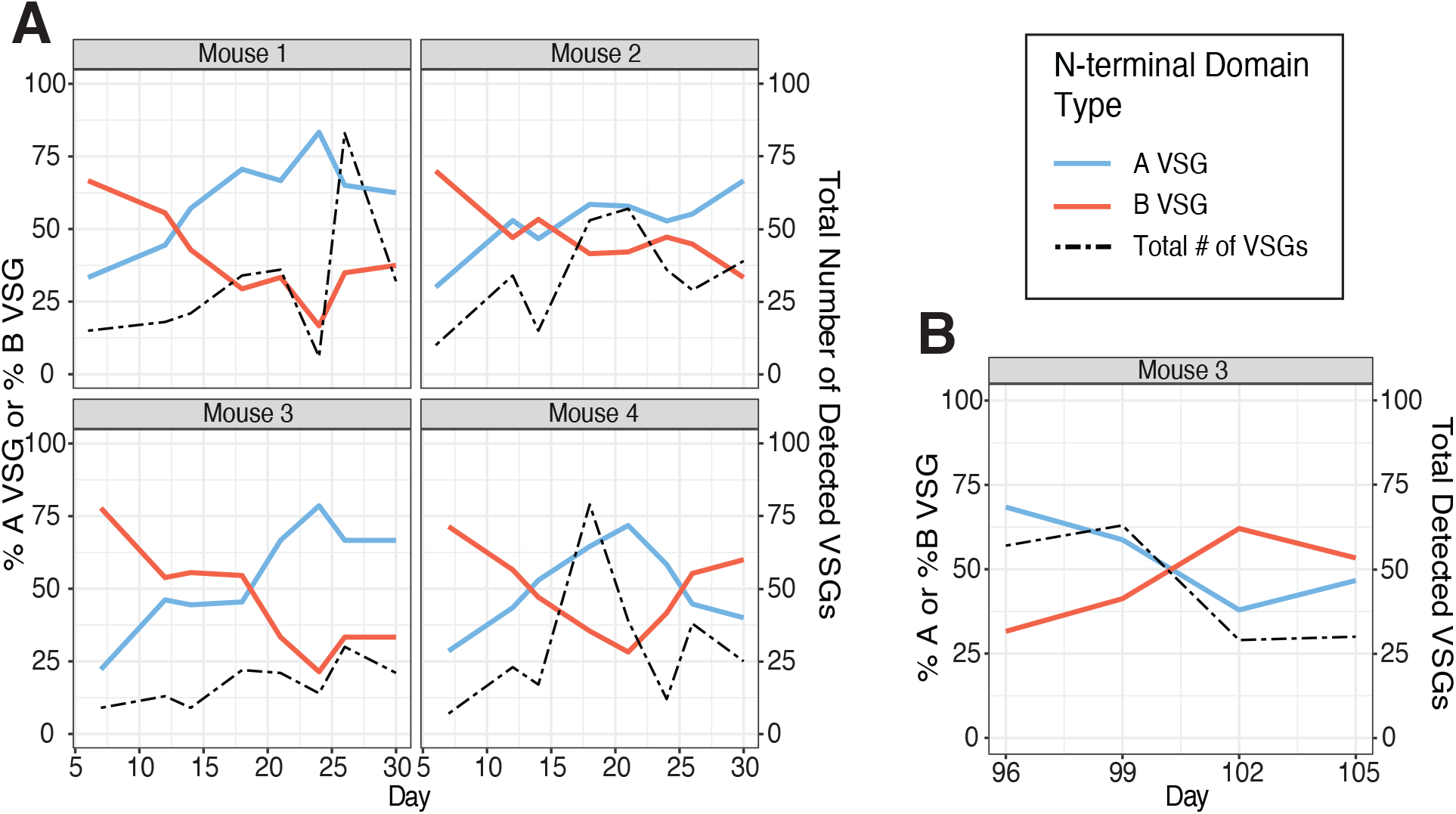
VSG N-terminal type composition fluctuates over the course of infection in mice. Proportions of N-terminal domain types expressed in *T. b. brucei* infected mice over time. The black dotted line represents the total number of identified VSGs. A) N-terminal type composition days 0-30. B) Type composition days 96-105.

**Supplemental Figure 8.**
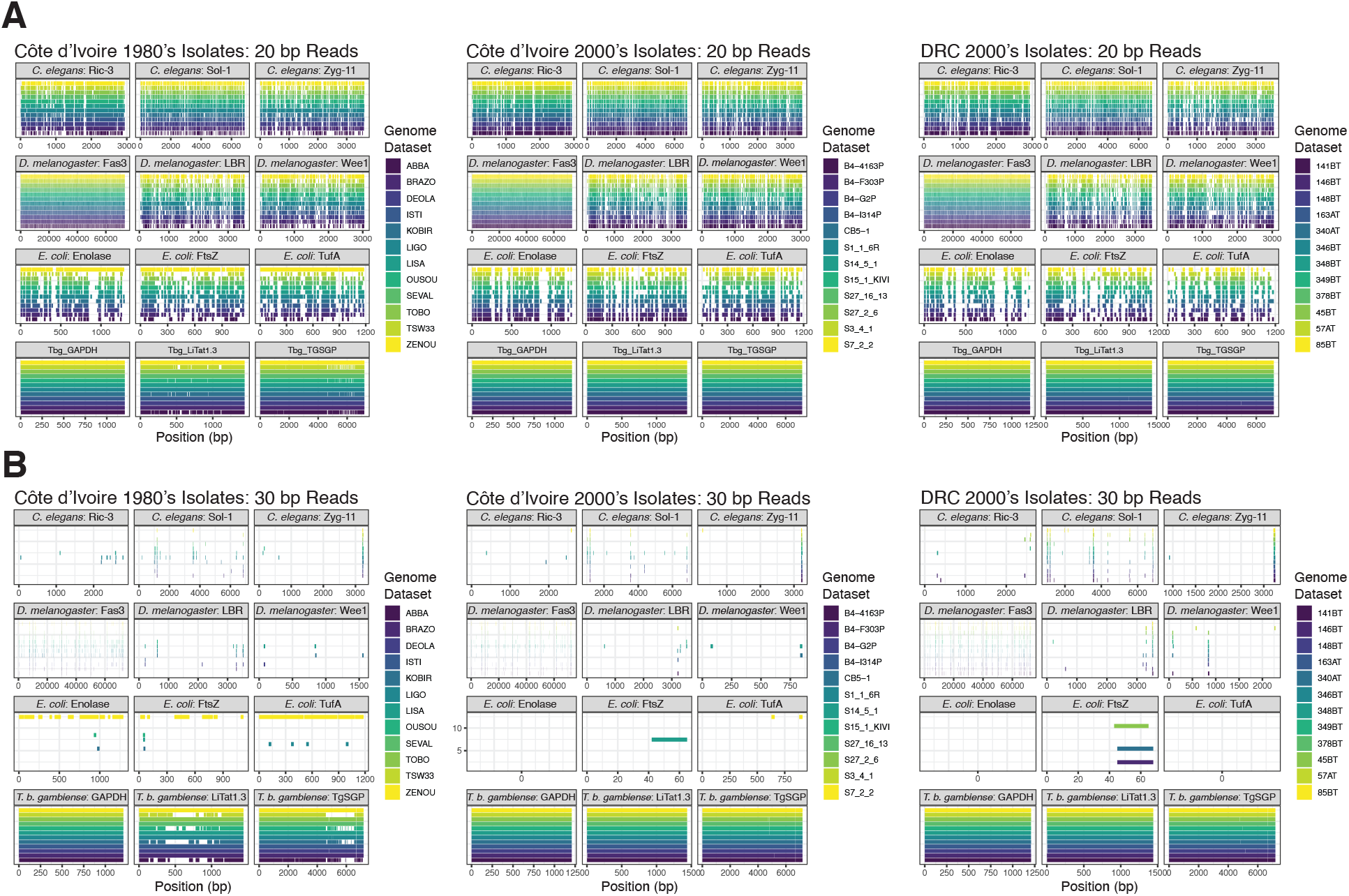
Mapping controls show how read size affects stringency of alignments, and support presence of sequences within datasets. A) Base pair coordinates of bowtie alignment ranges using 20 bp read lengths and allowing 2 mismatches for each of the 36 whole genome datasets. Positions of alignment hits are shown on the x-axis and each facet shows results for the 9 negative controls as well as 3 *T. b. gambiense* gene positive controls. The negative controls are randomly selected genes from other model organisms. B) Base pair coordinates for the same set of positive and negative gene mapping controls using 30 bp read lengths and allowing 2 mismatches. Coverage of the negative control genes is greatly reduced, while the *T. b. gambiense* gene positive controls still have alignment hits across the entirety of the gene.

**Supplemental Figure 9.**
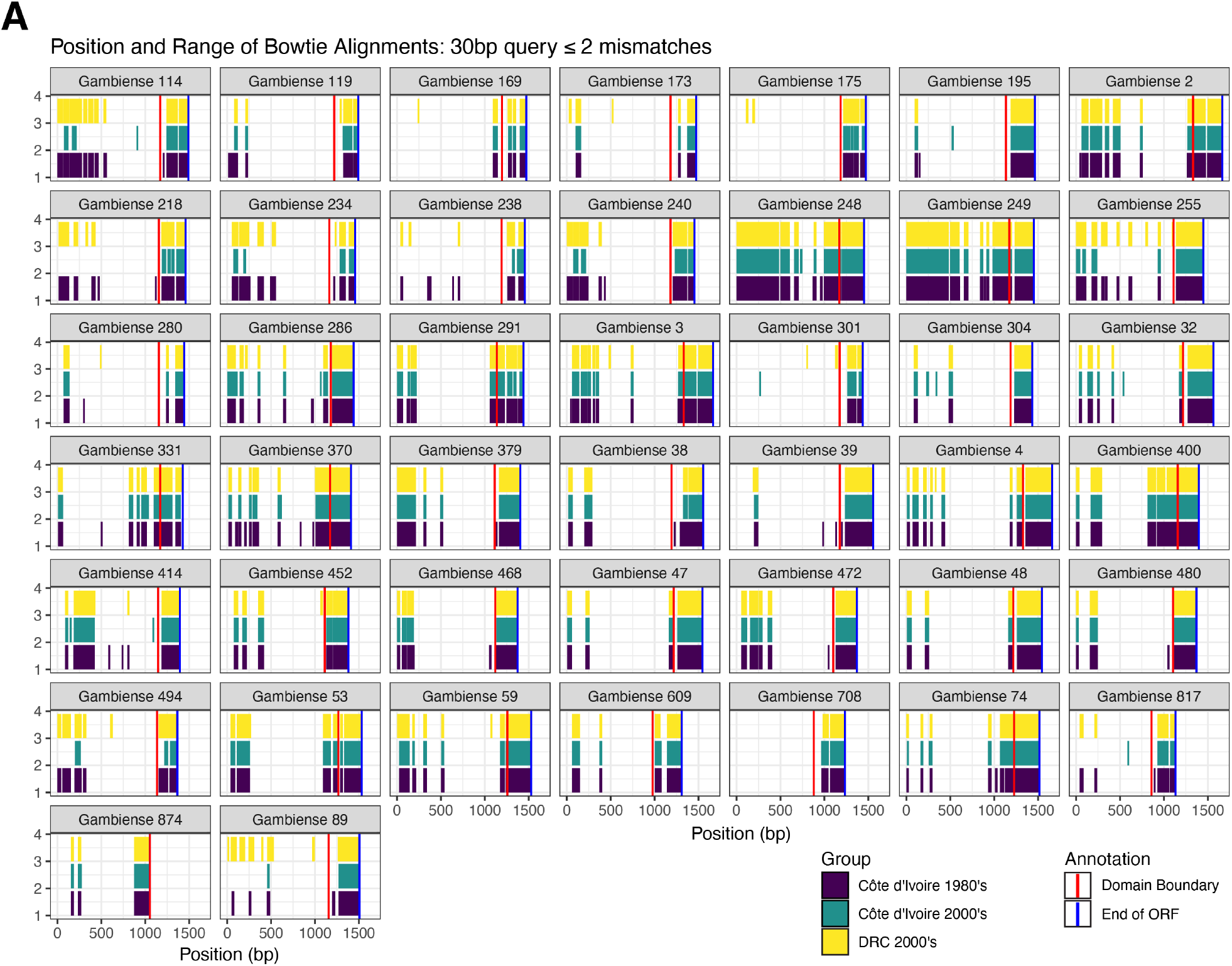
Summary of Bowtie alignment hits for each assembled gHAT patient VSG against the genomic sequences. A) Base-pair coordinates of each patient VSG are plotted as the X-axis, and each facet designates the patient VSG as well as the full ORF sequence length. Bars color-coded by genome dataset group show alignment length and position within the VSG ORF sequence for genomic sequence fragments of 30bp in length.

