## Supplementary figures and images for "VSGs expressed during natural *T. b. gambiense* infection exhibit extensive sequence divergence and a subspecies-specific expression bias"

### Supplemental Figure 4. BLASTp-tree of all T. b. gambiense VSGs. File attached.

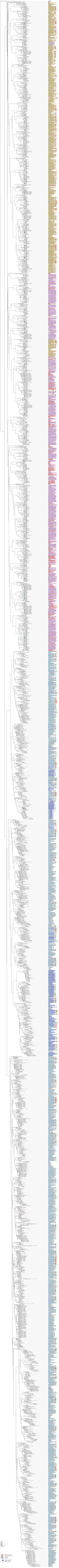
